# Molecular mechanisms of spontaneous curvature and softening in complex lipid bilayer mixtures

**DOI:** 10.1101/2022.02.17.480963

**Authors:** Henry J. Lessen, Kayla C. Sapp, Andrew H. Beaven, Rana Ashkar, Alexander J. Sodt

**Affiliations:** Eunice Kennedy Shriver National Institutes of Child Health and Human Development, National Institutes of Health, Rockville, MD 20892*; Postdoctoral Research Associate Program, National Institute of General Medical Sciences, National Institutes of Health, Rockville, MD 20892; Department of Physics, Virginia Tech, Blacksburg, VA 24061; Center for Soft Matter and Biological Physics, Virginia Tech, Blacksburg, VA 24061

## Abstract

Membrane reshaping is an essential biological process. The chemical composition of lipid membranes determines their mechanical properties, and thus the energetics of their shape. Hundreds of distinct lipid species make up native bilayers, and this diversity complicates efforts to uncover what compositional factors drive membrane stability in cells. Simplifying assumptions, therefore, are used to generate quantitative predictions of bilayer dynamics based on lipid composition. One assumption commonly used is that “per lipid” mechanical properties are both additive and constant — that they are an intrinsic property of lipids independent of the surrounding composition. Related to this, is the assumption that lipid bulkiness, or “shape” determines its curvature preference, independently of context. In this study, all-atom molecular dynamics simulations on three separate multi-lipid systems were used to explicitly test these assumptions, applying methodology recently developed to isolate properties of single lipids or nanometer-scale patches of lipids. The curvature preference of populations of lipid conformations were inferred from their redistribution on a dynamically fluctuating bilayer. Representative populations were extracted by both structural similarity and semi-automated hidden Markov model analysis. The curvature preferences of lipid dimers were then determined and compared to an additive model that combines the monomer curvature preference of both the individual lipids. In all three systems, we identified conformational subpopulations of lipid dimers that showed non-additive curvature preference, in each case mediated by a special chemical interaction (e.g., hydrogen bonding). Our study highlights the importance of specific chemical interactions between lipids in multicomponent bilayers and the impact of interactions on bilayer stiffness. We identify two mechanisms of bilayer softening: *Diffusional softening*, which is driven by the dynamic coupling between lipid distributions and membrane undulations, and *conformational softening*, which is driven by the inter-conversion between distinct dimeric conformations.

## INTRODUCTION

Compositional heterogeneity in cellular membranes is necessary for function [1]. The bilayer and its constituent lipids are important for many essential functions including: signaling [2–4], chemical separation [5], intracellular sorting [6], metabolism [7], and biogenesis [8]. In eukaryotic cells, different organelles have well defined lipid compositions [1] and deviations from normal lipid composition indicate stress or pathology. Membranes also show compositional asymmetry in their leaflets [9], enriching the possibilities for functional roles of lipid composition. Finally, bilayers are not well mixed within the leaflet and frequently display enrichment of particular lipid types in specific regions and shapes [10, 11]. For example, lipid-dependent spatial-correlations of surface receptors directly ties compositional affinity to cellular signaling processes [12]. Stresses due to lipid chemistry (i.e. deviation from preferred geometry) not only influence local mechanics but also protein function [13–15]. Because of the extensive physiological role and great chemical diversity of lipids in the cell, it is imperative to understand how the energetics of native bilayers depend on its many components.

More work is required to relate findings from model membranes (typically one to three lipid species) to biological membranes (hundreds of lipid species). It is often assumed that membrane properties are directly proportional to relative amounts of different lipids, but this is not always the case. For example, cell-extracted giant plasma membrane vesicles have lower bending modulii (*κ*_b_) than model counterparts. The bending modulus of cell-extracted giant plasma membrane vesicles was measured to be ca. 20 *k*_B_*T* [16]. This is considerably lower than membranes of fluid membranes like DOPC (26 *k*_B_*T* [17]) and a liquid disordered phase mixture of DOPC, cholesterol, and sphingomyelin (46 *k*_B_*T* [18]). The same observation has been seen when comparing the area compressibility modulus of red blood cell extracted vesicles to model membrane mixture representative of the major lipid types [19]. The bending stiffness is of fundamental interest to membrane remodeling events, where it controls the overall energetic scale. The mechanism behind *κ*_b_ reduction is important for understanding the forces that drive many cellular processes.

To address the discrepancies between biological systems and their simplified models, it is necessary to understand how lipid mixtures differ from homogeneous bilayers. Leibler has previously described how bilayers will soften as a result of the dynamic redistribution of curvature-sensitive inclusions [20]. In the Supporting Information we derive (using the Helfrich/Canham model modified for local lipid energetics [21]) the softening factor *α* yielding the apparent bending modulus *κ*_apparent_ in our own notation:

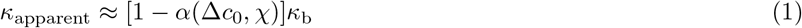

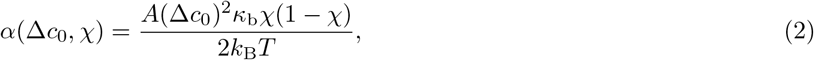

where Δ*c*_0_ is the difference of spontaneous curvature of the inclusion compared to the bulk, *χ* is the inclusion mole fraction, *κ*_b_ is the bilayer bending modulus, and *A* is the area of the inclusion. Consider, for example, a mixture of PE and PC lipids, with Δ*c*_0_ ≈ 0.02 Å^−1^, *A* ≈ 65 Å^2^, χ = 0.5, and *κ*_b_ ≈ 14 kcal/mol yields an 8% reduction in the bending modulus.

The correct identification of both the temporal and spatial scale driving softening is necessary for identifying the softening mechanism. As such, we have defined two possible mechanisms of softening that rely on differences in the spontaneous curvature of lipids: *diffusional* and *conformational*. Our proposed mechanisms differ in two major ways: origin of Δ*c*_0_ and timescale of redistribution. *Diffusional* softening is caused by the lateral distribution of lipids with different curvature preferences to membrane undulations. The timescale for this softening depends on both the timescales of lipid diffusion and the membrane undulations [21]. For example, a bilayer composed of minor species lipid A (non-zero curvature) and matrix lipid B (zero curvature) would exhibit this softening. Alternatively, *conformational* softening is due to the dynamic inter-conversion between curvature sensitive lipid conformations. This mechanism depends on the timescale governing inter-conversion between these curvature sensitive states. Importantly, knowledge of curvature-dependent lipid conformations is needed for this mechanism and is at odds with some common assumptions used in continuum membrane models.

Continuum models, like the Helfrich/Canham formalism [22, 23], use simplifying assumptions to calculate the energetics of membrane deformations. One such assumption is that per lipid continuum properties (i.e. spontaneous curvature, bending modulus, and area compressibility) are additive; i.e. the properties of the whole system are the linear sum of its parts. Employing the additive assumption dramatically reduces the complexity of heterogeneous membrane energetics, and indeed the assumption is supported experimentally in many cases. For example, the spontaneous curvature of cholesterol is similar in both PE and PC matrices [24]. This is, however, not always the case. Experiments on hexagonal phases indicate non-additive curvature preferences of palmitoyl sphingomyelin (PSM) mixtures with certain ceramides [25]. In cholesterol-rich bilayers, simulations indicate curvature energetics vary non-linearly with tail saturation [26]. Additionally, the formation of specific molecular interactions (in this case, hydrogen bonds) leads to a non-linear relationship between lipid content and curvature preference [26]. These exceptions highlight the importance of understanding how complex formation between lipid components alters the additive assumption for the continuum description of the bilayer. Current molecular simulation methods provide unparalleled resolution for identifying the molecular underpinnings of bilayer energetics. To take advantage of these simulations, new tools must be generated to efficiently identify novel lipid pairs and analyze their per lipid continuum properties.

To accomplish this, we developed a novel analysis that uncovers the properties of collections of lipids, including dimers. The analysis takes two approaches to isolating important dimer structures. The first directly identifies dimers that contained features suspected to be strong, such as salt-bridged interactions and hydrogen bonding. The second, a hidden Markov model (HMM), allows for the emergence of structural features based on their similarity in space and time. We consider this approach to be “semi-automated” in that the important lipid sites were chosen (e.g., charged moieties, hydrogen bond donors) as structural information but were not enforced to map to states.

Markov models have shown promise in being able to extract mechanistic insight and timescale information from molecular simulations. The Markov model framework provides a formalism for defining timescales for processes [27]. It typically relies on a discrete assignment of state such that transitions between states can be counted. They have been successfully employed to study several problems in molecular biophysics, including protein folding [28–32], ligand binding [33], protein-protein associations [34], and phase identification in lipid bilayers [35].

*Hidden* Markov Models (HMMs) allow for a weak association of an observable and the state of interest. In an HMM, there is not a one-to-one mapping of an observation to state. Rather, observed configurations are assigned to a state based on the time sequence of observations. Applications to lipid bilayers are particularly amenable to HMM analysis. First, the large number of target molecules in a simulation is directly proportional to the sampling statistics that can be obtained through simulation. In simulation systems containing hundreds of lipid molecules, even somewhat rare states can be sampled. Second, the molecular environment around a lipid relaxes much more slowly than for material solvated by water. This strongly suggests a hidden contextual influence on lipid configuration.

In this work, we use three unique test cases to examine varied effects of lipid-lipid interaction on the preferred curvature (spontaneous curvature) of lipid dimers. We introduce the additive model as the null hypothesis to which we compare dimer curvature preference. Sub-conformations of lipid dimers that exhibit non-additive spontaneous curvatures are then identified as curvature-sensitive states. In all cases, curvature-sensitive states were most easily identified by standard interaction metrics (e.g. hydrogen bond formation and divalent cation binding). These were predominantly distance based cutoffs for chemical sites known to form relatively strong interactions. The three systems studied with the approach were: 1) a tertiary mixture of 1,2-dioleoyl-sn-glycero-3-phosphocholine (DOPC), 1,2-dioleoyl-sn-glycero-3-phosphoethanolamine (DOPE), and 1,2-dioleoyl-sn-glycero-3-phospho-L-serine (DOPS), 2) a mixture of phosphatidyl-inositol-4,5-bisphosphate (PIP2) and 1-palmitoyl-2-oleoyl-glycero-3-phosphocholine (POPC), 3) binary mixtures of PSM and POPC. These three mixtures highlight three unique mechanisms that can alter the spontaneous curvature of a dimer conformation. Specifically, the DOPC, DOPE, and DOPS system illustrates the impact of hydrogen bond formation on the negative spontaneous curvature of PE. The more physiologically relevant systems (2 and 3) help to provide mechanistic insights into lipid-lipid interactions that are important for cellular function. We observed that Ca^2+^-meditated hetero-dimers between PIP2 and POPC have highly negative spontaneous curvature. The implication is that one powerful curvature-generating mechanism for PIP2 does not require clustering. Additionally, the PSM and POPC mixtures reveal a complicated role for hydrogen bonding that alters dimer spontaneous curvature in a way that appears to depend on the lipid matrix. As PSM is a major component in the plasma membrane and has been observed in nanometer-scale domains in cells [36], this finding has potentially broad-reaching implication for a number of membrane trafficking processes that depend on membrane mechanics.

## METHODS

### Molecular Dynamics

#### Build and simulation parameters

All simulations used the CHARMM C36 lipid forcefield [37]. Both PC/PE/PS and PSM/POPC simulations were built to be oblong, with one lateral dimension four times larger than the other, to capture long wavelength undulations. Simulations of PIP2/POPC were built square. Four replicas of PC/PE/PS each had 1272 total lipids. One replica each of 10, 20, or 30% PSM in POPC had 1320 total lipids. One simulation of 10:1 PIP2:POPC had total 484 lipids, with 129 Ca^2+^ions. When simulating PIP2, the CHARMM residues SAPI24 and SAPI25 were used in equal amounts. The residues only vary by the protonation state of either the fourth or fifth position phosphate group. These lipids both possess one stearoyl (18 carbon, fully saturated) and one arachidonic (20 carbon, 4 unsaturated bonds) acyl tail each. All systems were simulated for at least one microsecond (summing all replicas) with a constant temperature of 310.15 K and pressure of 1 bar. A summary of components and durations for each of the simulation systems is listed in Table S2 of the Supporting Information.

Using NAMD [38], systems were minimized and initially relaxed steps as prescribed by CHARMM-GUI. After these steps, the systems were converted into AMBER format [39, 40] using ParmEd. All analyzed data was simulated with AMBER. Constant temperature was targeted and regulated by a Langevin thermostat with a friction coefficient of 1 ps^−1^, and constant pressure anisotropically (*x* and *y* coupled; zero surface tension) maintained by a Monte Carlo barostat. Non-bonded interactions were switched off between 10–12 Å. Long-range electrostatics were handled by the PME implementation for GPUs by Salomon-Ferrer et al. [41]. The PME grid was constructed with a spacing of less than 1 Å (tolerance = 10^−5^; Ewald coefficient = 0.22664; interpolation order = 4). Bond lengths involving hydrogen were constrained by SHAKE [42]; water was kept rigid using SETTLE [43]. A 2 fs time step was used and coordinates were saved every 200 ps for analysis.

### Dimer identification

Lipid structures were characterized by a subsets of atoms, defined for each type below. Pairs of lipids were selected if the center-of-geometry of the subset was within 14 Å. Atom names are listed below in the C36 lipid forcefield naming scheme.

#### Hydrogen bonding from DOPE

The atom selections for dioleoyl (DO) phosphatidylethanolamine (DOPE), DO-phosphatidylcholine (PC, DOPC) and DO-phosphatidylserine (PS, DOPS) were `{P, N, C1, C3, C24`, and `C34}`. Hydrogen bonds between lipids were identified when a PE hydrogen was within 2.5 Å of a hydrogen bond acceptor, here either the phosphate oxygens or the carbonyl oxygens.

#### Divalent-cation mediation of PIP2

Both SAPI24 and SAPI25 PIP2 residues were treated identically in lipid pair detection. The atom selections for PIP2 were {C11, C12, C13, C14, C15, C16, P4, P5, P, C1, C2, C3, C21, C31}, while POPC used `{P, N, C1, C11}`. Divalentcation mediated dimers were identified when a Ca^2+^ (or Mg^2+^, see Supplemental Material) ion were within 4 Å of the center-of-geometry of a lipid dimer.

#### Hydrogen-bonding of PSM

The atom selections for PSM were `{N, P, NF, HNF, O3, HO3`, and `OF}`. Hydrogen bonds between lipids were identified when either the `HNF` or `HO3` atom was within 2.5 Å of a hydrogen bond acceptor, here a phosphate oxygen, `OF`, or `O3`.

#### General state identification with an HMM

The full description of the HMM methodology for dimer state extraction is described in the Supplemental Material. Briefly, a subset of dimer states were extracted using the atom selections and cutoffs above, yielding between 10^4^ and 2 × 10^4^ structures for a dimer type. Fifty-two clusters were formed by K-Medoid clustering, which chooses clusters based on representative structures from a set, associating each member of the whole set with a cluster center to minimize the summed RMSD of the clusters. These clusters formed the observable in the HMM. Following the definition of the clusters, all time-ordered sequences of dimers were extracted from the simulation, assigning each instance of a dimer to a cluster based on RMSD. An HMM describing the observed sequences of dimer clusters was created by optimizing the transition and observation probabilities of the HMM such that the probability of observing the ordered sequences was maximized. State was then assigned to each dimer configuration based on the optimized HMM.

### Computation of dimer curvature preference

We define here the curvature-coupled redistribution (CCR) method, which extracts the curvature experienced by lipids, implying the spontaneous curvature [21]. A cartoon illustrating the overall approach is shown in Fig. 1.

**FIG. 1:**
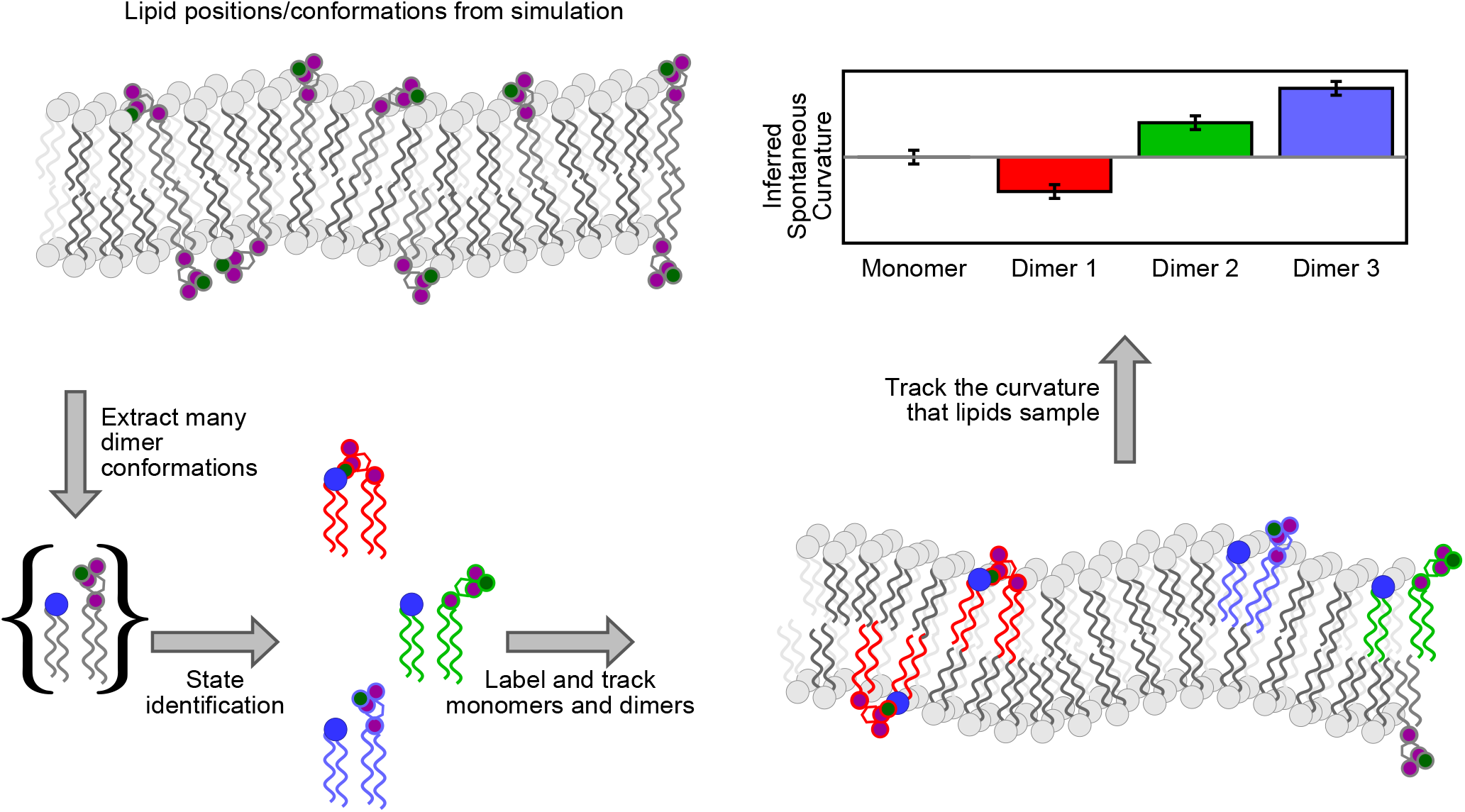
A cartoon illustrating the curvature-coupled redistribution (CCR) method. Beginning at the top left and proceeding counter-clockwise, a patch of multi-component bilayer is simulated (represented above by purple and gray lipids in the top left). A set of dimer conformations are extracted and collected into states with common properties. Here, the cartoon represents divalent-cation mediated interactions between an anionic lipid and its neighbors. The correlation of state (i.e., molecular structure) and curvature is recorded and compared with Helfrich/Canham theory. The mechanism of curvature sensitivity of a state is inferred from the correlation of structure and sampled curvature.

The time-resolved discrete Fourier spectrum is computed on a grid with maximum spacing 15 Å. The height is computed first for each leaflet; the bilayer height is the mean of the two. The coefficient *h*_q_ of mode **q** is

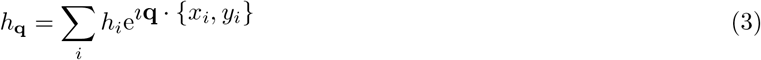

where here *h_i_* is the height of the grid point located at lateral coordinates {*x_i_*, *y_i_*} For each time frame, the set {*h*_q_} is recorded, along with the lateral ({*x*, *y*}) positions for each lipid.

The time-resolved curvature a lipid experiences is computed as ∇^2^*h*(*x, y*):

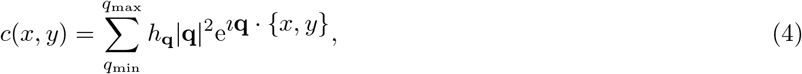

which is real as 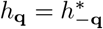. The upper cutoff, *q*_max_ determines the highest frequency mode used to compute curvature. The *q*_max_ used in our analysis is 0.08. The average curvature experienced by a lipid is then the average of *c*(*x, y*) over the trajectory.

#### The additive model for lipid dimers

A simple hypothesis for the spontaneous curvature of dimers is that their spontaneous curvature is the area-weighted average of the two species, and that the total area of the dimer is the sum of the monomer areas:

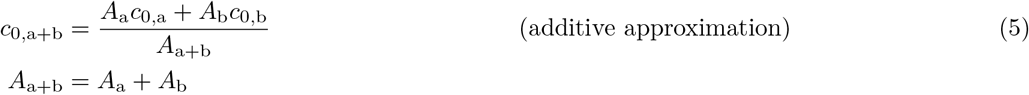

From the Helfrich/Canham model [22, 23] and assuming a localized lipid mechanical extent [21], the expected curvature of some embedded molecule x on mode *q* is proportional to its area fraction (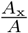, where *A*_x_ is the area of the embedded object and *A* is the area of the bilayer patch) times its spontaneous curvature difference with the background, Δ*c*_0_ [21]:

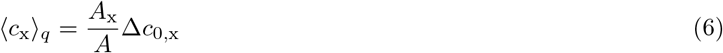

The *additive model* of two interacting lipids is that the expected curvature for a dimer is the sum of the observed monomer curvatures:

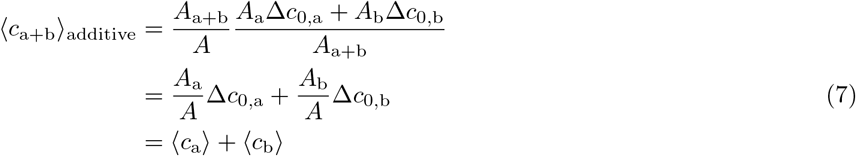

Consider the random co-localization of two PE lipids under the additive approximation. As a single unit, the patch will have twice the influence on curvature compared with a single lipid, all else being equal.

## RESULTS

### The additive model as the null hypothesis

Our study is primarily concerned with determining the impact of complex formation on dimer curvature preference. We will therefore investigate all three of the test systems described above using the same relative reference state: the additive model. In this model there is no effect of specific interactions between neighboring lipids on curvature. The Δ*c*_0_ used to construct the model comes from the bulk lipid redistribution analysis and are reported as the single lipid species differences in spontaneous curvature. Values of Δ*c*_0_ for each monomer in this paper are reported in Table I. Use Eq. 7 to translate between Δ*c*_0_ and experienced curvature, 〈*c*〉 (as reported in Figs. 2, 3, and 4).

**TABLE I:** Δ*c*_0_ values used in the additive model. These Δ*c*_0_ values are calculated from a bulk analysis of spontaneous curvature values for individual lipid species subtracted from the spontaneous curvature of the entire membrane.

| Lipid | $\Delta c_0$ ($\text{\AA}^{-1}$) | Simulation |
| --- | --- | --- |
| DOPE | $-0.015 \pm 0.002$ | 30 % DOPE 25% DOPC +<br>45% DOPS |
| DOPC | $+0.010 \pm 0.002$ | |
| DOPS | $+0.003 \pm 0.001$ | |
| PIP2 | $-0.028 \pm 0.006$ | 10% PIP2 90% POPC |
| POPC | $+0.004 \pm 0.001$ | |
| PSM | $-0.001 \pm 0.002$ | 10% PSM 90% POPC |
| POPC | $+0.003 \pm 0.0002$ | |
| PSM | $+0.001 \pm 0.0009$ | 20% PSM 80% POPC |
| POPC | $-0.0002 \pm 0.0002$ | |
| PSM | $+0.003 \pm 0.0006$ | 30% PSM 70% POPC |
| POPC | $-0.0005 \pm 0.0002$ | |

**FIG. 2:**
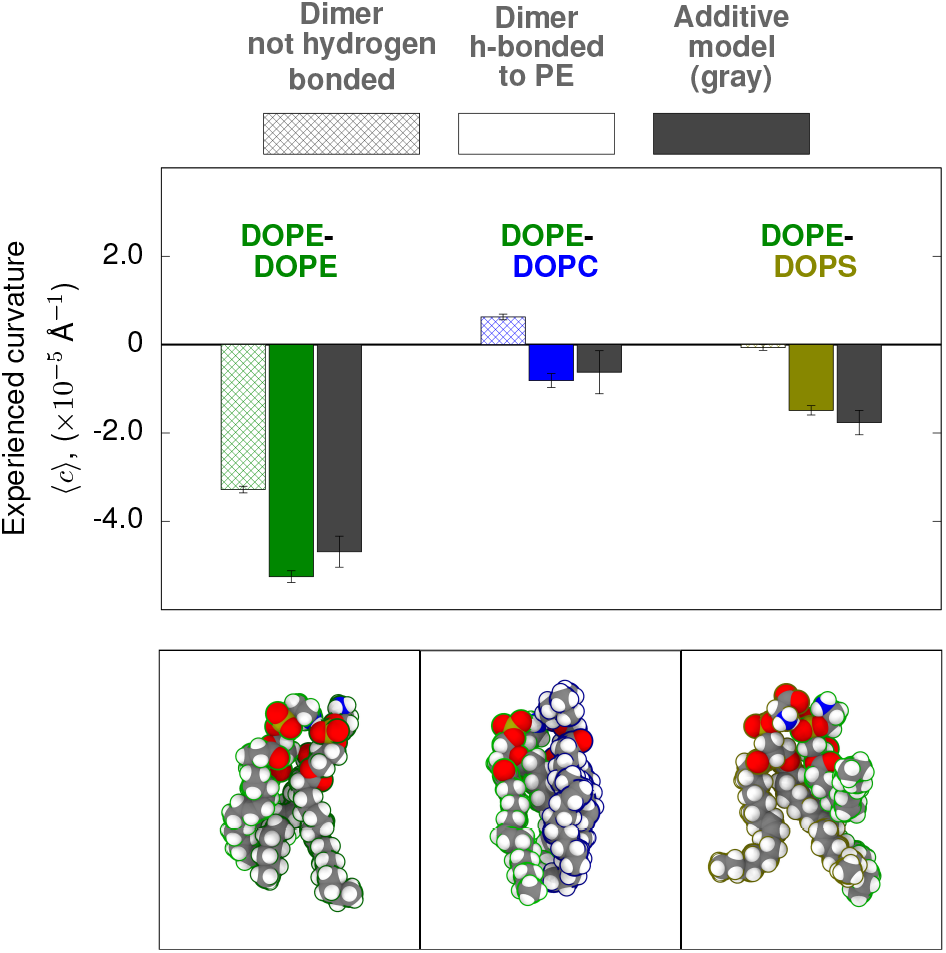
Hydrogen-bond donation from PE headgroups enhances negative curvature in lipid dimers. Legend above figure (in gray) shows the mesh bars represent the non-hydrogen bonded complexes, the colored bars represent the hydrogen bonded dimers, and the gray bar represents the theoretical, additive model. Experienced curvature is plotted for lipids as monomers or as in complex. Lipids (DOPE, green; DOPC, blue; DOPS, gold) are colored by type. The additive model is shown in gray. The bottom panels are representative molecular configurations of hydrogen bonded dimers.

**FIG. 3:**
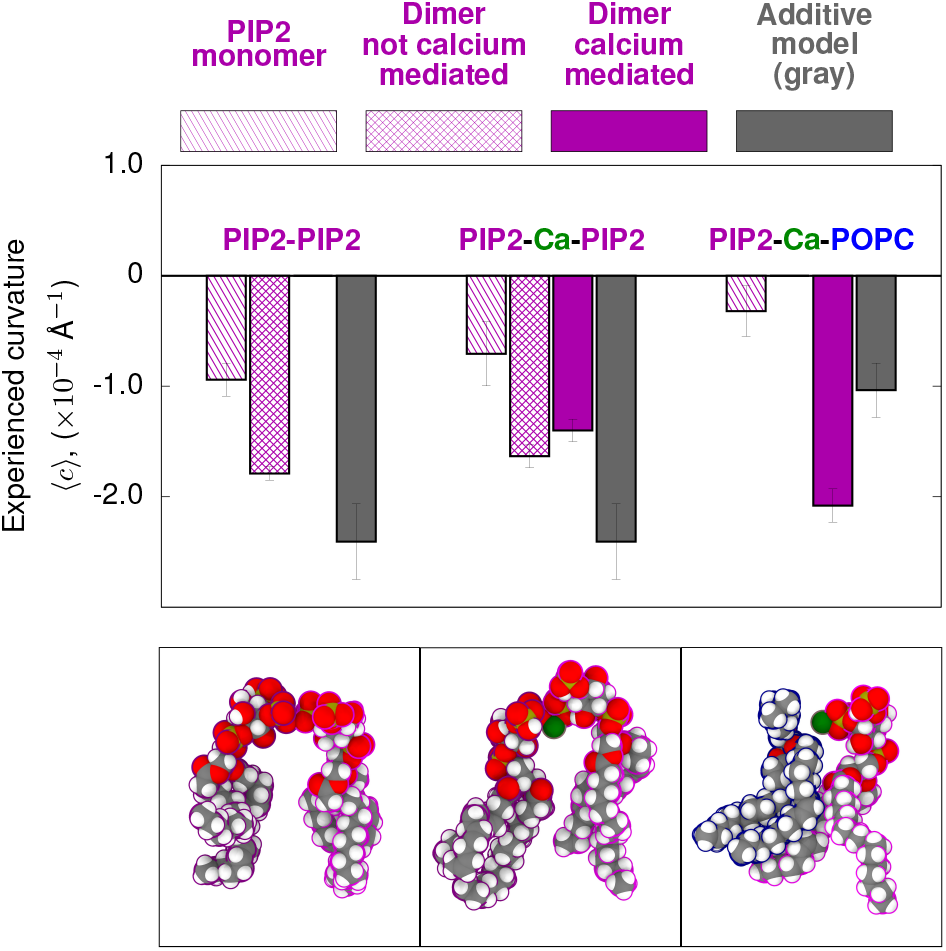
Calcium bridged interactions between PIP2 and POPC show strong negative curvature. Experienced curvature is plotted for lipids as monomers or as in complex. The additive model is shown in gray, monomers are striped. Dimers are meshed. Ca^2+^-mediated dimers are solid. The bottom panels are representative molecular configurations of each type (PIP2-PIP2, PIP2-Ca^2+^-PIP2, and PIP2-Ca^2+^-POPC).

**FIG. 4:**
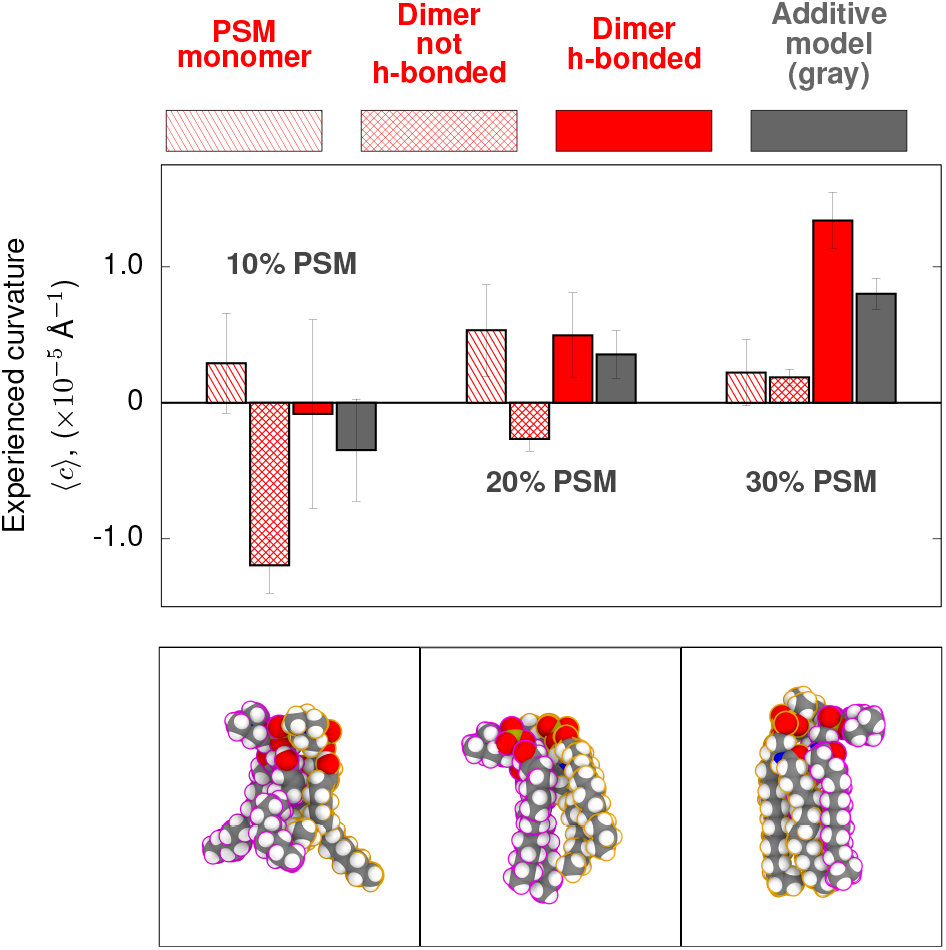
Curvature sensitivity of PSM dimers changes with hydrogen-bond formation and concentration. Experienced curvature is plotted for lipids as monomers or as in complex. The additive model is shown in gray, monomers are striped. Dimers are meshed. Hydrogen-bond-mediated dimers are solid. The bottom panels are representative molecular configurations of the amide-hydrogen-bonded configuration.

Dimer curvature preferences predicted by the additive model are then compared to observed curvature preferences measured for lipid dimers. In Figs. 2, 3, and 4, monomer curvature is shown by bars patterned with stripes, dimers are shown with mesh patterns, and interaction-mediated dimers (hydrogen bonding, calcium-bridged) are shown as solid colored bars. Expected values according to the additive model are shown with solid gray bars. Significant deviation from the additive model by the complex indicates a breakdown of the assumption of monomer independence. In all three systems, states exhibiting non-additive curvature preference were identified. We use the correlation of structural features and curvature to infer molecular mechanisms for non-additive dimer spontaneous curvature.

Monomers are defined as those lipids that do not meet the criteria for a dimer. For example, when classifying PIP2-PIP2 dimers, PIP2 lipids as members of PIP2-Ca^2+^-POPC complexes would be classified as monomers if they are not in complex with other PIP2 lipids. When identifying PIP2-Ca^2+^-POPC complexes, PIP2 lipids as members of PIP2-PIP2 complexes would be classified as monomers if they are not bridged by Ca^2+^ to POPC.

### Hydrogen bond donation from DOPE lipids induces negative curvature

Figure 2 reports the average curvature experienced by dimers of DOPE with DOPE, DOPC, or DOPS. The simulations in this work have been previously used in Ref. [21] to establish the equivalence between the lateral pressure profile and CCR methodology. It is clear that DOPE modeled with the CHARMM C36 forcefield has a negative spontaneous curvature, consistent with experiment [24]. Also apparent from this analysis is the observation that complex formation with DOPE leads to significant deviations from the additive approximation.

The spontaneous curvature of lipid dimers containing DOPE depends on variations of the intra-complex conformations of interacting lipids. Two mechanisms are proposed to influence the spontaneous curvature of DOPE: headgroup size and hydrogen bond formation [44–46]. The small PE headgroup suggests it is a predominant factor in determining its negative spontaneous curvature. The present analysis indicates, however, this is not the complete story. A comparison of dimers in which DOPE forms a hydrogen bond (middle column, solid in Fig. 2) or does not (left column, hashed) indicates substantially increased negative spontaneous curvature for configurations with hydrogen bonding. This highlights the importance of specific molecular interactions when relating lipid content to curvature.

Following theory (Eq. 1), this system should undergo both diffusional and conformational softening. DOPE can form complexes between all three types of lipids in the system, each with a different bulk curvature preference. The dynamic, diffusional coupling of these complexes to membrane undulations will lower the apparent bending modulus. Likewise, formation of a hydrogen bond in each of these complexes significantly changes co to become more negative. Rapid inter-conversion between these states then leads to a softer bilayer. Even in a system with only 3 components, lipid heterogeneity and molecular details complicate the traditional continuum interpretation of bilayer mechanics.

### Divalent-cation mediated interactions of PIP2 induce negative curvature independent of clustering

Three PIP2 dimer schemes were analyzed. In the first, a dimer is defined by its interactions with neighboring PIP2 (left column, Fig. 3). Redistribution analysis indicated dimers were consistent with an additive model (compare gray and mesh bars). Thus, PIP2-PIP2 interactions did not appear to influence curvature. Second, PIP2-PIP2 dimers were isolated if their interaction was mediated by Ca^2+^ (center column, Fig. 3). Again, no significant negative curvature influence of the interaction was detected, compared with the additive model. Third, the dimer state was defined as a Ca^2+^-mediated interaction with neighboring POPC (right column, Fig. 3). Here, a significant, non-additive negative curvature preference was detected.

These results support current models of PIP2 cluster curvature preference and highlight the need to better understand the role of divalent cations in mediating PIP2 heterodimer formation. While in native membranes PIP2 is a minor lipid species [47], can form clusters that enhance local concentration [48]. In this case, the PIP2 clusters should act as a slowly diffusing curvature-sensitive inclusion that contributes to softening. While the PIP2-PIP2 dimers within these clusters show a reduced negative curvature when compared to the additive approximation, these cluster are also surrounded by a myriad of other lipids. Our results suggest that further analysis of PIP2 clustering would show that curvature is affected by the boundary of a PIP2 domain, where conformations would more closely resemble the PIP2-Ca^2+^-POPC interaction we identify. These types of ion-mediated heterodimers present a potentially important case for conformational softening near PIP2 clusters that may have functional relevance for curvature influencing proteins that bind PIP2 [4].

### PSM complex formation promotes positive curvature

Figure 4 shows that the monomeric state of PSM exhibits similar curvature preferences to POPC (little net curvature preference). Neighboring PSM lipids *without* hydrogen bonds do not show a positive curvature preference (meshed bars). By comparison, PSM lipids that form amide-mediated hydrogen-bonded states exhibit a positive spontaneous curvature (Fig. 4 solid bars). This shift to a more positive spontaneous curvature becomes more pronounced as the mole fraction of PSM is increased. The relationship between the increased positive curvature preference of the hydrogen bonded state and the mol fraction of PSM suggests that formation of the amide hydrogen bond helps to drive shift to more positive spontaneous curvature at higher concentrations of PSM.

It is important to highlight that hydrogen bond formation in the PSM systems has the opposite effect (i.e. biasing the complex to more positive curvatures) that what was seen in the DOPE system (more negative curvature bias). The positioning of the PSM amide hydrogen bond within the leaflet is likely important for determining the sign of the curvature. Hydrogen bond formation occurs closer to the bilayer mid-plane in the PSM lipids than the hydrogen bonds made by lipids PE headgroups. In the case of DOPE, attractive forces near the headgroup surface lead to negative curvature [49]. We propose that amide HB formation observed in the PSM system helps to reduce tail entropy and, therefore, shift the curvature preference more positive (further explained in the Discussion).

PSM exhibits a concentration dependent positive curvature preference, yet it it is unclear that the amide-hydrogen-bonded dimer is sufficient to completely explain the tend. We speculate that the increasing positive curvature of PSM is also due to higher order effects beyond dimerization. For example, with sufficient PSM nearby, cohesive hydrogen bonding supports chain ordering, further lessening entropic tail repulsion and thus its contribution to positive curvature. A similar effect was observed with the chain-ordering ability of cholesterol [26]. The possibilities for extending the CCR analysis to higher order complexes are discussed below.

This system shows an additional way that heterogeneous lipid composition can complicate the traditional continuum view of mixed bilayers. In this case, changes in the lipid matrix composition influence both the sign of different PSM-PSM complex curvature preference and the magnitude of hydrogen bond formation on the curvature. By making the difference in curvature preferences between the hydrogen bonded and non-hydrogen bonded states larger, the anticipated magnitude of conformational softening will increase. Further, as matrix PSM increases, PSM-PSM dimers prefer more positive curvature, which changes where these structures will diffuse to and will alter the impact of diffusional softening.

## DISCUSSION

### Curvature preference is a balance of repulsive and cohesive forces along the bilayer normal

Curvature preference is typically described using the heuristic of “lipid shape” that conveniently describes lipids as cones or cylinders [50]. Needless to say, specific interactions between lipids are incompatible with shape. Thus, identification of the curvature sensitivity of specific interactions coupling to curvature refutes the ubiquity of the shape model. In contrast, CCR provides the required framework for distinguishing the molecular mechanisms of spontaneous curvature quantitatively.

It is the balance of repulsive and attractive forces that determines the curvature stresses of a leaflet, and thus its spontaneous curvature. Qualitatively, repulsive forces like occlusion (due to bulky head groups) and tail entropy are the essence of the shape model. Attractive forces, such as hydrogen bonding and Coulombic interactions, are better described by specific interactions. Both are quantified in the lateral pressure profiles [49] that are typically used to infer *c*_0_. The lateral pressure profile of PE, for example, has a cohesive stress at the depth of the hydrogen bond [51], relative to PC. There is currently no method to determine, from the pressure profile, whether this is due to the size of the headgroup or hydrogen bonding.

Applying the CCR method allows us to correlate molecular interactions with curvature, and make connections to the transverse positions of interactions. In the cases of both DOPE and Ca^2+^-mediated interactions of PIP2, attractive interactions near the level of the glycerophospholipid phosphate induce strong negative curvature. In contrast, the amide hydrogen bond is formed deeper in the membrane for PSM-PSM dimers. By anchoring the two lipids closer to the tails, we speculate that the hydrogen bond restricts the tail entropy and, therefore, reduces the repulsive forces on that region of the lipid. Lessening the tail repulsion would then give rise to the positive curvature preference of these PSM dimers. Additionally, positive curvature coincides with ordering of the surrounding matrix. That is, c0 is more positive at 30% PSM and ordered configurations are likely.

### Curvature-dependent lipid configurations determine the bending modulus

The bending modulus *κ*_m_ is the elastic constant describing the second-order variation in the curvature energy. The magnitude of *κ*_m_ is related to the conformational plasticity of the lipids. An example of this is the stiffening (ca. 5 times [52]) of ordered phase bilayers compared to disordered bilayers. In the liquid disordered phase, lipids are less tightly packed and can adopt a greater range of conformations, presumably leading to a smaller bending modulus. Conversely, the liquid ordered phase restricts lipid dynamics and increases packing, increasing the bending modulus. The differences between these two phases highlights the importance of molecular details when interpreting a bulk phenomenon like bending energy. We relate two mechanisms of bilayer softening that are displayed on very different timescales. First, consider *diffusional softening*, the change in the undulation relaxation spectrum due to dynamic lateral fluctuations of a mixture of lipids with varied c0. The second mechanism shows how membrane bending rigidity can be reduced through the interconversion between lipid pairs with distinct, curvature-dependent states, which we refer to as *conformational softening*.

#### Diffusional softening

A bilayer is assembled with symmetrical leaflet composition: a small molecular fraction χ of lipid A with spontaneous curvature *c*_a_ amidst a background of lipids each with zero spontaneous curvature. Further assume that there is a well-defined bending modulus *κ*_b_ that is the same for the background as well as that of lipid A. The bending modulus 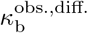 of the entire mixture will appear to be

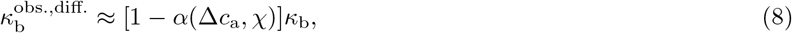

with a defined in Eq. 2. See the Supplemental Information for the derivation of Eq. 8. Here Δ*c*_a_ is the difference of spontaneous curvature between *c*_a_ and the spontaneous curvature of the background lipid composition. The softening factor (*α*(*c*_a_, χ)) arises from the coupling of the lateral distribution of lipids to bilayer undulations. Note that the lateral distribution of lipids relaxes slowly (as *D*^−1^*q*^−2^, where *D* is the diffusion constant and 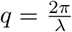 where λ is the undulation wavelength) for sub-micron wavelength undulations. This introduces kinetics as an important consideration when investigating softening mechanisms.

#### Conformational softening

Consider now softening due to (perhaps rapidly) interchanging curvature-dependent conformations. Our data shows that different conformations can exhibit different curvature preferences. We propose a simple model in which mechanical properties (spontaneous curvature and bending modulus) are altered by adding in a specific interaction. The hydrogen bond donated by a PE lipid to a neighbor adds negative curvature preference that appears to be largely independent of the acceptor (Fig. 2). The Ca^2+^-mediated interaction between PIP2 and POPC dramatically shifts curvature to be negative (Fig. 3). We model the interacting state as being “added on” by shifting the spontaneous curvature of a reference lipid, such as POPC, by *δ*. The reference lipid has area-per-lipid *A*_R_, spontaneous curvature *c*_0,R_ and bending modulus *κ*_b_. The likelihood of the interacting state is further characterized by its curvature-independent mole fraction, χ.

Apply separate local Helfrich models *H*_ref_. and *H*_int_. for the curvature energy of the reference and interacting states, respectively:

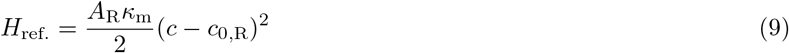

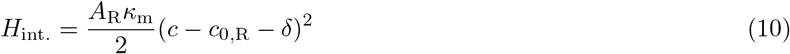

where *A*_R_ is the area per lipid for the reference, *c* is the curvature local to the lipid, and *κ*_m_ is the monolayer bending modulus. The free energy (*F*) is:

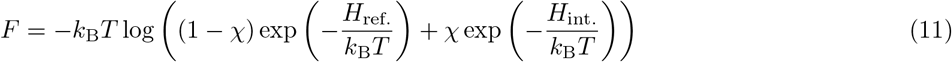

*F* can be restructured in a form resembling the Helfrich Hamiltonian. Consider the case in which *δ* is small. Expanding to second order in *δ*:

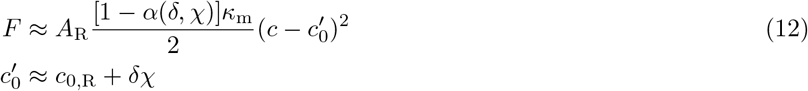

Where *α*(*δ*, χ) is the same as in Eq. 2. With the same factor *α*(*δ*, χ), compositional and conformational heterogeneity have a similar effect on bilayer properties. For *δ* = 0.03 Å^−1^, *A_p_* = 65 Å^−2^, χ = 0.5 and *κ*_m_ = 7 kcal/mol, the lipid patch has 83% the stiffness. This is a substantial decrease in the apparent stiffness of the bilayer, and can be directly attributed to the ability of specific states to stabilize different curvatures.

Unlike chemical composition, the conformational composition fluctuates rapidly. Furthermore, the interchange of conformations is typically much faster than the lateral rearrangement of lipids. An intuitive example is conformational isomerization of acyl chains, which occurs on the sub-nanosecond timescale [53], whereas lipids require 10s of nanoseconds to change neighbors, and *D*^−1^*q*^−2^ for a fluctuation with wavenumber *q* to relax (microseconds for a tens of nanometers lengthscale).

Molecular configurations of lipids are a continuum, not discrete. The reasoning in this section thus only forms a rationale for how curvature sensitivity of conformational sets determines softening, rather than for predicting *κ*_b_ *ab initio*. We have implicitly chosen *κ*_b_ to represent the bending modulus of some idealized simple lipid, for example, POPC, that serves as a base value. *Stiffening*, on the other hand may result in the *removal* of conformational space, for which the base of *κ*_b_ of POPC, and the treatment here, would be inadequate.

Equilibrium measurements that report the entire distribution of undulation amplitudes (diffusive X-ray studies and fluctuation analysis at low *q*) will show the effects of both diffusion and conformational softening. However, neutron spin echo (NSE) experiments of lipid mixtures will be dominated by the fast relaxation processes that determine conformational softening, and would likely be insensitive to diffusion softening. Consider for example the NSE experiments on mixtures of DOPC and cholesterol in Ref. [54] which find that a DOPC bilayer is stiffened by cholesterol. Diffusional softening is expected by cholesterol, a lipid that induces high negative curvature [24], yet this would only be clear from techniques able to probe long timescales in which lateral fluctuations of cholesterol concentration are relaxed. This is a plausible explanation for the disagreement between diffusive X-ray studies and NSE techniques [55, 56]. Combining the two techniques would isolate mechanisms of softening that are important at separate timescales.

### Impact on bilayer viscosity

The undulation of a simple bilayer relaxes with rate 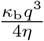 at low *q*. For higher frequency undulations, inter-leaflet friction and surface viscosity slow the rate [57]. Relaxation times at short length and timescales contribute to the characterization of membrane mechanics by NSE. Experimental work using homogeneous PC lipid bilayers suggest that acyl chain dynamics are important determinants of viscosity [58]. Extending insights drawn from reduced model systems [59], the authors speculate that both internal dynamics controlled by factors such as tail saturation and external factors (e.g. lipid-lipid coupling) control different timescale relaxation phenomena [58].

In addition to acyl chain dynamics, ensembles of curvature-sensitive conformations must also relax; we hypothesize that the lipids studied herein, as well as others with similar chemistry, should contribute to the viscosity. The lifetimes of complexes studied here (many nanoseconds for PIP2-Ca^2+^-POPC bridging) are on the same order of magnitudes for the slow relaxation times reported in studies of alcohol and alkane systems [59] and DMPC bilayers [58]. Plausibly, long-lived lipid dimers would increase the lipid-lipid couplings that are proposed to dominate at higher *q*. In this way, we identify two competing ways that viscosity can be altered by headgroup dynamics and heterogeneity in multi-component bilayers (diffusional and conformational). In lieu of the additional molecular mechanism implicated in determining bilayer viscosity, new whole lipid approaches are needed for investigating the viscoelastic properties of multi-component bilayers.

### Mechanics is a likely target of lipid homeostasis and compositional complexity

The unique chemistry of PIP2 allows it to play an outsized role in healthy cell physiology. Despite being 1% of cellular membrane composition [60], it is an essential part of cellular pathways including actin dynamics, cell adhesion, endocytosis, signaling, and organelle trafficking [61, 62]. In these processes, PIP2 binds various effector proteins that can either be peripheral membrane proteins (e.g. BAR proteins, kinases, etc), or integral membrane proteins (e.g. voltage gated potassium channels). PIP2 function is tied to cellular localization that depends intimately on the physical chemistry of the lipid. It is a negatively charged lipid due to the addition of phosphate groups, and as a result can make a strong complex with divalent cations, but only if the cation is present at sufficient concentration [63].

Divalent cation binding is important to PIP2 membrane biophysics. Studies have shown that PIP2 organization in lipid bilayers is altered by the presence of divalent cations [64]. Namely, PIP2 is suggested to form micro-domains within the native membrane and thereby increase its effective local concentration [64]. While calcium is a physiologically important ion, clustering also is induced by magnesium and zinc [48]. Computational studies have used graph theory and network analysis to describe the differences these various ions have on interconnectivity induced by these ions [65]. Simulations of PIP2 indicate it has negative spontaneous curvature, with divalent cation binding increasing the coupling to negative curvature [66]. This adds an additional layer of complexity to understanding micro-domain formation in PIP2. In addition to knowing that PIP2 can exist at high effective concentrations, it is also important to consider the local bilayer structure of PIP2 micro-domains.

In Ref. [66], the negative curvature of Ca^2+^-bound PIP2 is studied in pure PIP2 bilayers. The authors found that high calcium concentrations (2:1 Ca^2+^:PIP2) are necessary to induce the strongest negative curvature. In contrast, we find that at 10% coverage, it is the simple Ca^2+^-mediated interactions with neighboring POPC that induce negative curvature, an effect that would not be apparent in a bilayer saturated with PIP2. According to our analysis, *domains* of PIP2 are not required for a profound mechanical effect. Rather, it is the strong connection to neighboring lipids by divalent cations that is required.

## CONCLUSIONS

This study introduces methodology for linking molecular interactions to lipid spontaneous curvature. The negative spontaneous curvature of PE lipids is partly explained by its ability to donate hydrogen bonds to neighboring lipids. Divalent-cation mediated interactions between PIP2 and neighboring glycerophospholipids show a strong trend to negative curvature, indicating that domains of PIP2 are not required for its negative curvature preference. In a third case, hydrogen-bond linked conformations of PSM showed a strong positive curvature preference only at higher PSM concentrations, suggesting the ordered lipid matrix is part of the mechanism.

The shape of membranes is typically interpreted in terms of the average, or monomer, mechanical properties of lipids, assuming static or uniform distributions of lipids. Biological membranes, however, are neither static nor uniform. By ignoring the possibility of substates with different curvature preferences or averaging over these substates, it is likely that important mechanistic insights into physiological process are being missed. The simulation methodology here presents a path to understanding this complexity, with the mechanism testable by scattering and membrane-fluctuation analysis techniques. In our judgment, the modeling of *κ*_b_ with diffusional and conformational softening, the characterization of the spatial mechanical extent of lipids and multi-lipid complexes [21], and the understanding of the effects of differential stresses between leaflets [67], is forging a clear path to characterization of the bending modulus and curvature stresses of biological membranes with complex composition.

## ACKNOWLEDGMENTS

This work was supported by the intramural research program of the *Eunice Kennedy Shriver* National Institutes of Child Health and Human Development (NICHD) at the National Institutes of Health. A.H.B. was supported by a Postdoctoral Research Associate (PRAT) fellowship from the National Institute of General Medical Sciences (NIGMS), award number 1Fi2GM137844-01. Molecular rendering was performed with Tachyon software written by John E. Stone. Simulations were performed on computational resources provided by the NICHD.

