## Supplemental Material for "Molecular mechanisms of spontaneous curvature and softening in complex lipid bilayer mixtures"

### I. HMM STATE DETECTION

A general overview of the HMM methodology used in this study will be provided. First, dimer conformations are extracted from MD trajectories from user defined parameters that will be described in detail below. These structures are clustered and subsequently divided into HMM observables that describe the energy landscape. The HMM observables are then used to define hidden states and optimize parameters that define the model. The states are then used in additional calculations to determine curvature preference of specific dimer subpopulations from their dynamic redistribution (curvature-coupled redistribution, CCR). A graphical summary of the steps in the HMM/CCR method is shown in Figure 1, and a list of terms is presented in Table S1.

| Term | Meaning |
|---|---|
| **Approach complex** | A single instantaneous configuration of two lipids with centers separated by less than $R_{\mathrm{pair}} = 14$ Å. A limited set of atoms of the approach complex are used to compute mean-squared-displacements (MSDs) |
| **Similarity function** | The criteria for judging the similarity of two approach complexes by weighting RMSD and hydrogen-bonding patterns. These are labeled as **S**, with subscripts indicating function parameters. |
| **Similarity center** | A representative configuration from an ensemble of similar approach complexes. Determined by K-Medoid clustering [1]. The “center” of the cluster. |
| **Similarity cluster** | The set of approach complexes more similar to a particular similarity center than to any other similarity center. These are the observables of the HMM. |
| **Pair trajectory** | The time-ordered sequence of similarity cluster that is assigned to a pair of lipids. |
| **HMM state** | The target state of a lipid-lipid complex: A lipid pair possibly rich with structural and curvature-sensitive features, but which is identified only by the time sequence of its assigned similarity cluster. Determined by optimizing a HMM. The HMM state is computed by applying the HMM to a pair trajectory. |
| **Hidden Markov model** | The kinetic model of inter-conversion between hidden Markov states, and the probabilities of observing any HMM state as a member of a similarity cluster. |

TABLE S1: Terms used to describe the configurations and models for developing the HMM.

#### A. Using an HMM to extract states

For example, PSM approach complexes are identified using five headgroup atoms from each PSM molecule (10 atom positions from each dimer). The atoms are { `OF`, `C1F`, `HNF`, `O3`, `HO3` }. For a complete list of atoms used to define approach complexes in each system analyzed, refer to Table S1. These atom positions are used to calculate the distance between molecules using the center of geometry. Pairs of lipids within $R_{\mathrm{pair}} = 14$ Å were considered approach complexes. Atom positions are also used to calculate MSD between different approach complexes (used for scoring in K-medoid clustering).

We used approximately 15,000 approach complexes per simulation to define a pool from which the similarity centers were extracted. The set was extracted by traversing the trajectory uniformly, taking all complexes within $R_{\mathrm{pair}}$. The quantity was limited by increasing the amount of trajectory skipped. Our software was able to efficiently determine similarity centers when the pool was less than 20,000.

Additional considerations must be made when identifying ion-mediated approach complexes. When processing simulations to extract this class of approach complex, an ion type and distance cutoff must be specified in the processing input file. When searching for the bridging ion, first the approach complex center (based on the specified atoms) is calculated. Then the distances from this point and the ions proximal to the approach complex are calculated to find the closest ion. If distance from the ion is then compared to the maximum radial distance (4 Å), this ion is taken as the bridging ion.

##### *Construction of the HMM*

K-medoid clustering [1] was used to reduce the complexity of the extracted approach complexes and identify HMM observable states. This method uses both the mean-square-displacement (MSD) between approach complexes (following rotational and translational alignment) and a heuristic hydrogen bond matching scheme to rank similarity between approach complexes. The unweighted MSD between two approach complexes with a matching hydrogen bond were reduced by $\epsilon_+$ Å$^2$ for each bond, where $\epsilon_+$ is a tunable parameter. If a hydrogen bond was present

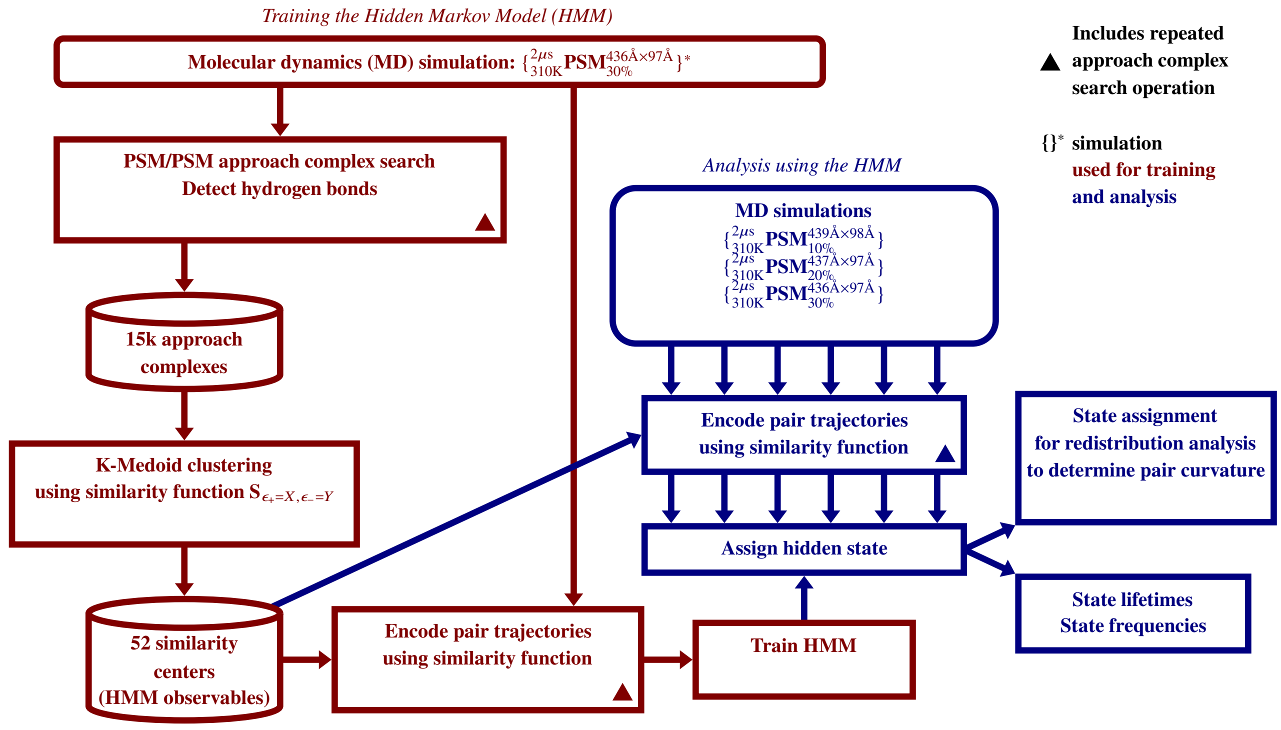


FIG. S1: A flowchart showing the process of dimer discovery and assignment via a hidden Markov model (HMM). The training (left side) and analysis (right side) steps are distinguished by color.

in one of the approach complexes but not the other, the MSD was penalized by $\epsilon_-$ Å$^2$. An exception was made for the amide-amide sphingolipid hydrogen bond. Penalties were applied that restricted members of a similarity cluster to have this bond (or the lack thereof) in common. The identity of one approach complex's **1** and **2** were swapped during the comparison, and the smaller MSD was selected, yielding a process independent of **1**/**2** ordering. The function applied by the K-medoid clustering to compare two pairs is called a similarity function, and any one function is denoted as **S**, with $\epsilon_\pm$ parameters indicated in the subscript. The final value of the similarity function is denoted $\sim\chi^2$. For each similarity function, the number of medoid centers was set to 52, defining *similarity centers* labeled { A-Z, a-z }.

The data was then processed to describe the time evolution of the approach complexes using the similarity centers determined through K-medoid clustering.This is done by reiterating through the MD trajectory in order to assign individual approach complexes to specific similarity clusters throughout the simulation. When processing the MD trajectory to build the HMM, the same spacing used for extracting approach complexes was used. An approach complex was assigned the letter code of the medoid center with which it had the lowest $\sim\chi^2$. This yields many distinct series of similarity cluster observables, stored as simple character strings. These distinct series (called a pair trajectory) describe the evolution of independent approach complexes. The pair trajectories are then used to build the kinetic portion of the HMM.

Finally, we have to determine the "hidden" states used in our HMM, which will will refer to as HMM states. The model defines $N_s = 6$ states. There is no correct number of states no more than there is a correct model for any physical system. Rather, if "too many" states are selected, an interesting state may be split into two states that interchange rapidly, but which exchange slowly with the other states. If "too few" states are selected, features of interest can not sort cleanly into states. Quotes here indicate that these are qualitative judgments. Once states are determined, all approach complexes are then classified using the HMM states and used for further analysis, such as the curvature preference calculations described below.

#### II. PIP2: PROTONATED ON THE 4- OR 5-PHOSPHATE

Figure S2 shows the curvature coupling of PIP2 protonated on the 4-phosphate (left) and the 5-phosphate (right). While there are differences between the two states, results for each protonation state are consistent with the obser-

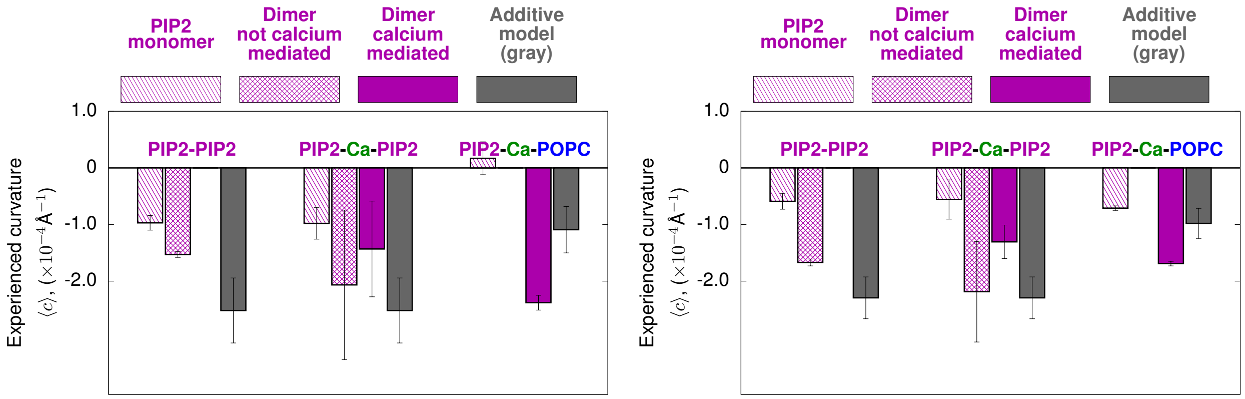


FIG. S2: PIP2 protonated on the 4-phosphate (left) and the 5-phosphate (right).

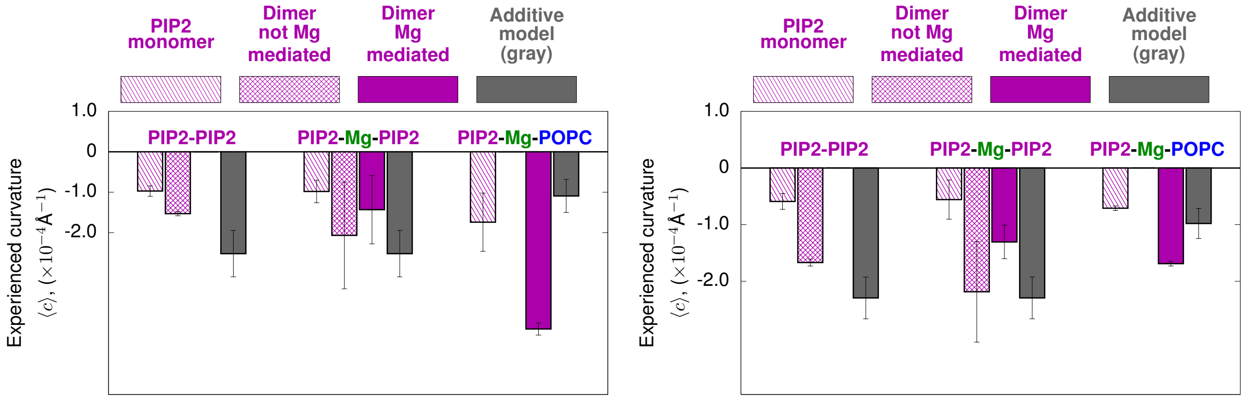


FIG. S3: PIP2 spontaneous curvature, but now with interactions bridged by $Mg^{2+}$instead of $Ca^{2+}$. PIP2 is protonated either on the 4-phosphate (left) or the 5-phosphate (right).

vation that PIP2-$Ca^{2+}$-POPC induces strong negative curvature.

#### III. PIP2: CALCIUM VS. MAGNESIUM IONS

Figure S3 shows the same curvature-dependent dimers now coupled by $Mg^{2+}$instead of $Ca^{2+}$. Results for $Mg^{2+}$are consistent with the mechanism that it is the cation-mediated interaction with neighboring generic phospholipid (here, POPC) that induces strong negative curvature.

#### IV. SOFTENING THEORY FOR COMPLEX BILAYERS

With

$$H_{\mathrm{HC}} = \frac{\kappa_{\mathrm{b}}}{2}(c_1 + c_2 - c_0)^2 + \bar{\kappa} c_1 c_2, \tag{S1}$$

the total elastic curvature energy is:

$$E_{\mathrm{HC}} = \frac{\kappa_{\mathrm{b}}}{2} \int_A \mathrm{d}\boldsymbol{S} H_{\mathrm{HC}}, \tag{S2}$$

To account for the preferred curvature $\Delta c_0 = c_{0,\mathrm{p}} - c_{0,\mathrm{background}}$ of a lipid ($c_{0,\mathrm{p}}$) in a background spontaneous curvature ($c_{0,\mathrm{background}}$), the energy density is modified by

$$H_\mathrm{c} = \frac{\kappa_\mathrm{m} A_p}{2} \int \mathrm{d}\boldsymbol{S} \big[ \left(c_1 + c_2 - c_{0,\mathrm{p}}\right)^2 - \left(c_1 + c_2 - c_{0,\mathrm{background}}\right)^2 \big], \tag{S3}$$

Here we do not include the Gaussian curvature modulus $\bar{\kappa}$ as we assume all lipids have equivalent $\bar{\kappa}$, and the integral of the Gaussian curvature does not vary with fluctuations.

For simplicity, the membrane surface, and thus its form in Eqs. S2 and S3, is parameterized in the linearized Monge gauge, a simplified surface parameterization capable of describing nearly planar configurations. The parameterization is $\boldsymbol{r}(x,y) = \{x, y, h(x,y)\}$ and linearization imposes a quadratic order cut-off [2, 3] that is accurate for thermal (here, weak) fluctuations:

$$E_\mathrm{H}\left[h(\boldsymbol{r})\right] = \int_A \mathrm{d}\boldsymbol{S}\, \frac{\kappa_\mathrm{b}}{2} \left[\boldsymbol{\nabla}^2 h(\boldsymbol{r})\right]^2 \tag{S4}$$

In Fourier space:

$$h(\boldsymbol{r}) = \frac{1}{A} \sum_{\boldsymbol{q}} h_{\boldsymbol{q}} \mathrm{e}^{\imath \boldsymbol{q} \cdot \boldsymbol{r}} \tag{S5}$$

$$h_{\boldsymbol{q}} = \int_A \mathrm{d}\boldsymbol{r} h(\boldsymbol{r}) \mathrm{e}^{-\imath \boldsymbol{q} \cdot \boldsymbol{r}}. \tag{S6}$$

For a real space function $f(\boldsymbol{r})$ the relation $f_{\boldsymbol{q}} = f^*_{-\boldsymbol{q}}$ applies to the Fourier transform coefficients. The coefficients are not independent. Furthermore, terms that must be real are real only if $\boldsymbol{q}$ and $-\boldsymbol{q}$ are considered simultaneously. In recognition of this we sum over $\boldsymbol{q}$ using the shorthand $\{\boldsymbol{q} > 0\}$,

$$\{\boldsymbol{q} > 0\} \equiv \{q_x, q_y\} \tag{S7}$$

such that

$$\begin{cases} 0 < q_x < q_\mathrm{max} & \text{for } q_y = 0 \\ -q_\mathrm{max} < q_x < q_\mathrm{max} & \text{for } q_y > 0 \end{cases},$$

yielding two independent functions, $f_{\boldsymbol{q},a}$ and $f_{\boldsymbol{q},b}$:

$$\begin{aligned} f_{\boldsymbol{q},a}(\boldsymbol{r}) &= 2\mathrm{Re}\, f_{\boldsymbol{q}} \cos(\boldsymbol{q} \cdot \boldsymbol{r}) \\ f_{\boldsymbol{q},b}(\boldsymbol{r}) &= 2\mathrm{Im}\, f_{\boldsymbol{q}} \sin(\boldsymbol{q} \cdot \boldsymbol{r}) \end{aligned} \tag{S8}$$

Henceforth we use the cos function's coefficient $f_{\boldsymbol{q},a} = 2\mathrm{Re}\, f_{\boldsymbol{q}}$ as the independent variable. Implicitly, the same derivations apply to the $b$-component, $h_{\boldsymbol{q},b}$.

The Fourier coefficients $\{\ h_{\boldsymbol{q},a}\ \}$ of the height (with $h_{\boldsymbol{q},b}$ defined analogously to $f_{\boldsymbol{q},b}$) are energetically uncoupled:

$$E_\mathrm{H}\left(\{h_{\boldsymbol{q},a}\}\right) = \frac{\kappa_\mathrm{b}}{2A} \sum_{\{\boldsymbol{q}>0\}} q^4 h^2_{\boldsymbol{q},a} \tag{S9}$$

with all other cross-terms energetically uncoupled at this order. Given a bilayer with a mole fraction $\chi$ of one lipid and $1-\chi$ background lipids, the energy of a density fluctuation is given by

$$\begin{aligned} E_\rho\left(\{p_{\boldsymbol{q},a}\}\right) &= \sum_{\{\boldsymbol{q}>0\}} \frac{k_\mathrm{B}T(p^2_{\boldsymbol{q},a})}{2A\rho_0\left(1-\chi\right)} \\ &= \sum_{\{\boldsymbol{q}>0\}} \frac{A_p k_\mathrm{B}T(p^2_{\boldsymbol{q},a})}{4A\chi\left(1-\chi\right)} \end{aligned} \tag{S10}$$

Where $p_{\boldsymbol{q},a}$ are Fourier coefficients for the distribution of particles (with $a$ and $b$ subscripts analogous to $f_{\boldsymbol{q},a}/f_{\boldsymbol{q},b}$ above), and $\rho_0 = \frac{2\chi}{A_p}$, with the factor of two representing the contribution from each leaflet. See derivation below.

The impact of the distribution of lipids with spontaneous curvature difference $\Delta c_{0,\mathrm{p}}$ is now introduced using the lipid density $\rho(\boldsymbol{r})$ which is convoluted with the spatial extent $w(\boldsymbol{r})$ to account for the change in curvature energy density in Eq. S3:

$$\begin{aligned}E_{\mathrm{c}}\left[h(\boldsymbol{r}),\rho(\boldsymbol{r})\right] &= \frac{\kappa_{\mathrm{m}}}{2}\int_{L^2}\mathrm{d}\boldsymbol{r}_p\int_{L^2}\mathrm{d}\boldsymbol{r}\rho(\boldsymbol{r}_p)w(\boldsymbol{r}-\boldsymbol{r}_p)\times\\ &\left[\left(\boldsymbol{\nabla}^2 h(\boldsymbol{r})-c_{0,\mathrm{p}}\right)^2-\left(\boldsymbol{\nabla}^2 h(\boldsymbol{r})\right)^2-c_{0,\mathrm{background}}\right]\\ &= \frac{A_p\kappa_{\mathrm{m}}}{2}\int_{L^2}\mathrm{d}\boldsymbol{r}\rho(\boldsymbol{r})\times\\ &\left[\left(\boldsymbol{\nabla}^2 h(\boldsymbol{r})-c_{0,\mathrm{p}}\right)^2-\left(\boldsymbol{\nabla}^2 h(\boldsymbol{r})\right)^2-c_{0,\mathrm{background}}\right]\end{aligned} \tag{S11}$$

Here Eq. S12 uses $w(\boldsymbol{r}-\boldsymbol{r}_p)=A_p\delta(\boldsymbol{r}-\boldsymbol{r}_p)$ as justified by the observed locality of PC, PE, and PS [4].

Performing the Fourier transform of Eq. S11 yields the particle-distribution/undulation coupling:

$$E_{\mathrm{c}}\left(\{h_{\boldsymbol{q},a}\},\{p_{\boldsymbol{q},a}\}\right) = -\sum_{\{\boldsymbol{q}>0\}}\frac{A_p\Delta c_{0,\mathrm{p}}\kappa_{\mathrm{m}}q^2 p_{\boldsymbol{q},a}h_{\boldsymbol{q},a}}{A}. \tag{S12}$$

with $p_{\boldsymbol{q},a}$ the Fourier coefficient for the density of lipid with spontaneous curvature differ $\Delta c_{0,\mathrm{p}}=c_{0,\mathrm{p}}-c_{0,\mathrm{background}}$. Combining Eqs. S9, S10, and S12, the total energy is

$$E_a = E_{\mathrm{H}}\left(\{h_{\boldsymbol{q},a}\}\right)+E_\rho\left(\{p_{\boldsymbol{q},a}\}\right)+E_{\mathrm{c}}\left(\{h_{\boldsymbol{q},a}\},\{p_{\boldsymbol{q},a}\}\right). \tag{S13}$$

The expectation of $h^2_{\boldsymbol{q},a}$ is determined by

$$\langle h^2_{\boldsymbol{q},a}\rangle = Z^{-1}\int \mathrm{d}h_{\boldsymbol{q},a}\mathrm{d}p_{\boldsymbol{q},a}h^2_{\boldsymbol{q},a}\mathrm{e}^{-\beta E_a}, \tag{S14}$$

where

$$Z=\int \mathrm{d}h_{\boldsymbol{q},a}\mathrm{d}p_{\boldsymbol{q},a}\mathrm{e}^{-\beta E_a}. \tag{S15}$$

For a single-component membrane ($\chi=0$ or $\Delta c_{0,\mathrm{p}}=0$), integration leads to

$$\langle h^2_{\boldsymbol{q},a}\rangle = \frac{Ak_{\mathrm{B}}T}{\kappa_{\mathrm{b}}q^4}\quad\text{(single component)}, \tag{S16}$$

from which the bending rigidity can be determined. For the two-component mixture, the expectation of $h^2_{\boldsymbol{q},a}$ is now:

$$\langle h^2_{\boldsymbol{q},a}\rangle = \frac{Ak_{\mathrm{B}}T}{\kappa_{\mathrm{b}}q^4}(1+\frac{\Delta c^2_{0,\mathrm{p}}A_p\kappa_{\mathrm{b}}\chi\left(1-\chi\right)}{2k_{\mathrm{B}}T}+\mathcal{O}[\Delta c^3_{0,\mathrm{p}}]) \tag{S17}$$

Thus, a bilayer with a symmetric or asymmetric mixture of lipids (with unequal spontaneous curvatures) will experience apparent softening according to

$$\begin{aligned}\kappa_{\mathrm{apparent}} &= \kappa_{\mathrm{b}}\left(1-\frac{\Delta c^2_{0,\mathrm{p}}A_p\kappa_{\mathrm{b}}\chi\left(1-\chi\right)}{2k_{\mathrm{B}}T}\right)\\ &= \kappa_{\mathrm{b}}\left(1-\alpha\right).\end{aligned} \tag{S18}$$

where $\alpha$ is the same as in Eq. 2 of the main text.

##### A. Fourier amplitudes of variations in lipid density

The derivation for the free-energy/entropic variation of the Fourier amplitudes of the density modes follows below.

Consider a bilayer with mol fraction $\chi$ of lipid A and $1-\chi$ of background lipids. Break the bilayer into $M$ narrow strips that are long in $y$ with length $L_y$ and narrow in $x$ with width $d$. There are $N$ lipids in each strip. The expected number of lipid $A$, $\langle n_A\rangle$ in each narrow strip is $\chi N$. Given random fluctuations, the variance of $n_A$ is $\chi(1-\chi)N$ according to the binomial distribution. For convenience define the density fluctuation function $\rho_A(i)$ as:

$$\Delta n_A(i) = n_A(i)-\chi N \tag{S19}$$

If the number of lipids in bin $i$, $n_A(i)$ is independent of $n_A(j)$, the autocorrelation function of $\Delta n_A(i)$, $r_{nn}(j-i)$ is

$$r_{nn}(j-i) = \begin{cases} \chi(1-\chi)N & : j = i \\ 0 & : j \neq i \end{cases} \tag{S20}$$

By the Wiener-Kninchin theorem, the power spectral density $S(f)$ is

$$\begin{aligned} S(f) &= \sum_{k=-\infty}^{\infty} r_{nn}(k)\mathrm{e}^{-i2\pi f k} \\ &= \chi(1-\chi)N, \end{aligned} \tag{S21}$$

that is, $S(f)$ is independent of $f$; it is "white noise." Given the power spectral density $S(f)$ in Eq. S21, discrete variables can be translated to the continuous case of interest: the Fourier transform of the lateral distribution of lipid $A$ density, $p_q$. Dividing by the bin area $dL_y$, transforming from $f$ to $q$ with $f = \frac{qL_x}{2\pi}$, and multiplying by 2 $\pi$ (by the convention in which the factor of $(2\pi)^{-1}$ is in the *inverse* transform) yields:

$$S(q) = (2\pi)\frac{L_x}{2\pi}\frac{1}{dL_y}S(f) \tag{S22}$$

$$= L_x \rho_A (1-\chi) \tag{S23}$$

Where here $\rho_A = \frac{N\chi}{dL_y}$ is the number of lipids per unit area. Instead of restricting analysis to $x$, performing the two-dimensional Fourier analysis with $\boldsymbol{q} = \{q_x, q_y\}$ yields

$$S(\boldsymbol{q}) = (1-\chi)A\rho_A \tag{S24}$$

Given a sufficiently large bilayer, we now substitute a normal distribution with matching variance. The probability $\mathrm{pr}(p_{\boldsymbol{q}})$ of observing density fluctuation $p_{\boldsymbol{q}}$ is given by

$$\mathrm{pr}(p_{\boldsymbol{q}}) \propto \exp\left(-\frac{p_{\boldsymbol{q},a}^2 + p_{\boldsymbol{q},b}^2}{(1-\chi)A\rho_A}\right) \tag{S25}$$

where $A_p$ is the area of the lipid, $p_{\boldsymbol{q},a}$ and $p_{\boldsymbol{q},b}$ are defined as for $h_{\boldsymbol{q},a}/h_{\boldsymbol{q},b}$ above. The expectation value $\langle p_{\boldsymbol{q}}^2 \rangle$:

$$\langle |p_{\boldsymbol{q}}|^2 \rangle = Z^{-1} \int_{-\infty}^{\infty}\int_{-\infty}^{\infty} \mathrm{d}p_{\boldsymbol{q},a}\mathrm{d}p_{\boldsymbol{q},b}(p_{\boldsymbol{q},a}^2 + p_{\boldsymbol{q},b}^2)\mathrm{e}^{-\frac{p_{\boldsymbol{q},a}^2 + p_{\boldsymbol{q},b}^2}{(1-\chi)A\rho_A}} \tag{S26}$$

$$= (1-\chi)A\rho_A \tag{S27}$$

with

$$Z = \int_{-\infty}^{\infty}\int_{-\infty}^{\infty} \mathrm{d}p_{\boldsymbol{q},a}\mathrm{d}p_{\boldsymbol{q},b}\mathrm{e}^{-\frac{p_{\boldsymbol{q},a}^2 + p_{\boldsymbol{q},b}^2}{(1-\chi)A\rho_A}} \tag{S28}$$

has yielded the correct variance.

#### V. TABULATED SIMULATION DETAILS


[1] Kaufman L. and Rousseeuw P. Clustering by means of Medoids. In *Stat. Data Anal. Based L1 Norm Relat. Methods*, pages 405–416, 1987.

[2] Frank L.H. Brown. Elastic modeling of biomembranes and lipid bilayers. *Annu. Rev. Phys. Chem.*, 59:685–712, 2008.

[3] Markus Deserno. Fluid lipid membranes: From differential geometry to curvature stresses. *Chem. Phys. Lipids*, 185:11–45, 2015.

[4] Kayla C. Sapp, Andrew H. Beaven, and Alexander J. Sodt. Spatial extent of a single lipid's influence on bilayer mechanics. *Phys. Rev. E*, 103(4), 2021.

| **System/composition** | Ions | Dimensions | Duration |
|---|---|---|---|
| 30% DOPE, 25% DOPC, 45% DOPS | 125 $Cl^-$ 697 $Na^+$ | 429×95×86 Å$^3$ | 4 × 600 ns |
| 10% PIP2 (half protonated on $P_4$, half on $P_5$) 90% POPC | 82 $Cl^-$ 129 $Ca^{2+}$ | 125×125×84 $\mathring{A}^3$ | 1×2 $\mu$s |
| 10% PIP2 (half protonated on $P_4$, half on $P_5$) 90% POPC | 82 $Mg^{2+}$ 129 $Ca^{2+}$ | 125×125×84 $\mathring{A}^3$ | 1×2 $\mu$s |
| 10% PSM 90% POPC | none | 432×98×95 Å$^3$ | 1×1.1 $\mu$s |
| 20% PSM 80% POPC | none | 423×98×97 Å$^3$ | 1×1.1 $\mu$s |
| 30% PSM 70% POPC | none | 415×98×98 Å$^3$ | 1×1.1 $\mu$s |

TABLE S2: Key descriptors of the simulations analyzed in this work.
